# Atypical relationships between neurofunctional features of print-sound integration and reading abilities in Chinese children with dyslexia

**DOI:** 10.1101/2021.11.11.468218

**Authors:** Zhichao Xia, Ting Yang, Xin Cui, Fumiko Hoeft, Hong Liu, Xianglin Zhang, Xiangping Liu, Hua Shu

**Affiliations:** State Key Laboratory of Cognitive Neuroscience and Learning & IDG/McGovern Institute for Brain Research, Beijing Normal University, China; School of Systems Science, Beijing Normal University, China; Faculty of Psychology, Beijing Normal University, China; Beijing Key Laboratory of Applied Experimental Psychology, National Demonstration Center for Experimental Psychology Education, Faculty of Psychology, Beijing Normal University, China; Haskins Laboratories, USA; Department of Psychological Sciences and Brain Imaging Research Center, University of Connecticut, USA; Department of Psychiatry and Weill Institute for Neurosciences, University of California, San Francisco, USA; Department of Neuropsychiatry, Keio University School of Medicine, Japan

**Keywords:** audiovisual integration, character, Chinese, dyslexia, individual differences, pinyin

## Abstract

Conquering print-sound mappings (e.g., grapheme-phoneme correspondence rules) is vital for developing fluent reading skills. In neuroimaging research, this ability can be indexed by activation differences between audiovisual congruent against incongruent conditions in brain areas such as the left superior temporal cortex. In line with it, individuals with dyslexia have difficulty in tasks requiring print-sound processing, accompanied by a reduced neural integration. However, existing evidence is almost restricted to alphabetic languages. Whether and how multisensory processing of print and sound is impaired in Chinese dyslexia remains underexplored. In this study, we applied a passive audiovisual integration paradigm with functional magnetic resonance imaging to investigate the possible dysfunctions in processing character-sound (opaque; semantics can be automatically accessed) and pinyin-sound associations (transparent; no particular meaning can be confirmed) in Chinese dyslexic children. Unexpectedly, the dyslexic group did not show reduced neural integration compared with typically developing readers in either character or pinyin experiment. However, the results revealed atypical correlations between neural integration and different reading abilities in dyslexia. Specifically, while the neural integration in the left inferior frontal cortex in processing character-sound pairs correlated with silent reading comprehension in both children with and without dyslexia, it was associated with morphological awareness (semantic-related) in controls but with rapid naming (phonological-related) in dyslexics. This result indicates Chinese dyslexic children may not use the same grapho-semantic processing strategy as their typical peers do. As for pinyin-sound processing, while a stronger neural integration in the direction of “congruent > incongruent” in the left occipito-temporal cortex and bilateral superior temporal cortices was associated with better oral reading fluency in the control group, an opposite pattern was found in dyslexia. This finding may reflect dyslexia’s dysfunctional recruitment of the regions in grapho-phonological processing, which further impedes character learning.

**Highlights:**

- Neurofunctional correlates of print-sound integration in Chinese children with and without dyslexia are investigated.
- Dyslexic children show atypical relationships between neural audiovisual integration and reading abilities.
- Chinese children with dyslexia are likely to use inefficient strategies to process characters and pinyin.

## 1 Introduction

Reading consists of multiple cognitive processes, and it takes years of formal instruction to achieve a high proficiency. In this process, establishing robust links between orthographic and phonological representations (e.g., grapheme-phoneme correspondence [GPC] rules) is one initial and fundamental step (Perfetti & Harris, 2013). Behavioral and neuroimaging studies of alphabetic languages reveal that it is critical to conquer the GPC rules to develop fluent reading skills. The failure will impede building efficient grapho-semantic mapping and eventually result in reading difficulties (Blomert, 2011; Di Folco, Guez, Peyre, & Ramus, 2020; Richlan, 2019; Shaywitz, 1998). Nowadays, most researchers agree that the manifestation of dyslexia is associated with linguistic features in a given language (Richlan, 2020). However, while existing evidence is almost restricted to alphabetic orthographies, the question of whether and to what extent print-sound integration is impaired in Chinese children with dyslexia remains underexplored, especially at the neurofunctional level.

Chinese has a morpheme-based logographic writing system (Perfetti, Cao, & Booth, 2013). In addition to phonological information, semantics is strongly involved in even the most fundamental processing—character recognition (Bi, Han, Weekes,& Shu, 2007; Guan, Fraundorf, & Perfetti, 2020; Liu et al., 2017; Yang, Shu, McCandliss, & Zevin, 2013; Zhao et al., 2014). At the behavioral level, longitudinal and meta-analytic studies have demonstrated the importance of both phonological-related (e.g., phonological awareness [PA], rapid naming [RAN]) and semantic-related (e.g., morphological awareness [MA]) skills in Chinese reading development (Pan et al., 2016; Lei et al., 2011; Liu et al., 2017; Ruan, Georgiou, Song, Li, & Shu, 2018). However, it should also be noted that the tasks used in these studies required explicit processing of the written scripts. A similar situation exists at the brain level. Previous functional magnetic resonance imaging (fMRI) studies revealed hypoactivation in the left inferior and middle frontal areas during visual rhythming and lexical decision in children with dyslexia, suggesting dysfunctions of the neural substrates underlying both print-to-sound and print-to-meaning mappings in tasks requiring explicit processing (Cao et al., 2016; Liu et al., 2012; Siok, Perfetti, Jin, & Tan, 2004). Paralleling the fMRI research, structural and diffusion imaging studies also provided evidence on alterations in morphometry of these regions and white matter tracts connecting them (Siok, Niu, Jin, Perfetti, & Tan, 2008; Su et al., 2018; Xia, Hoeft, Zhang, & Shu, 2016). Hence, while these findings indicate deficits in grapho-semantic and grapho-phonological processing in Chinese children with dyslexia, the question of whether implicit and automatic processing is impaired remains largely unknown.

In this study, we adopted a passive fMRI audiovisual paradigm (i.e., without explicit phonological or semantic judgment), which is appropriate for investigating automaticity in reading-related processing. This paradigm has been used in shallow orthographies such as Dutch and demonstrated the impaired letter-sound automatized integration as a likely proximal cause of dyslexia that is independent of phonological processing deficits (Blau et al., 2010; Blau, van Atteveldt, Ekkebus, Goebel, & Blomert, 2009; Blau, van Atteveldt, Formisano, Goebel, & Blomert, 2008). This paradigm’s basic logic is that if a brain area integrates auditory and visual inputs or is involved in the subsequent higher-level cognitive processes, its activation should differ between the congruent and incongruent conditions. This effect is usually named “congruency effect” when the activation in the congruent condition was stronger than the incongruent condition and is named “incongruency effect” otherwise. Here we used the term “audiovisual integration effect” (or “neural integration” for short), given that both directions indicate successful multimodal information integration. Since this effect can be observed even when no task or a passive task is used, researchers regard it to reflect implicit processing (Blau et al., 2010; van Atteveldt, Formisano, Goebel, & Blomert, 2007). To date, the neural integration has been demonstrated in skilled adult readers and typically developing children (Blau et al., 2010; van Atteveldt & Ansari, 2014; van Atteveldt, Formisano, Goebel, & Blomert, 2004; van Atteveldt, Formisano, Blomert, & Goebel, 2007).

Of importance, direction and strength of the neural integration are affected by several factors, such as characteristics of participants and orthographic depth of languages (Blau et al., 2010; Blau et al., 2009; Holloway, van Atteveldt, Blomert, & Ansari, 2015; Kronschnabel, Brem, Maurer, & Brandeis, 2014; Wang, Karipidis, Pleisch, Fraga-Gonzalez, & Brem, 2020). For example, individuals with dyslexia showed an atypical pattern in brain areas such as the superior temporal cortex (STC) (Blau et al., 2010; Blau et al., 2009). This anomaly was driven by hypo-activation in the congruent condition along with hyper-activation in the incongruent condition in dyslexia, indicating reduced neural integration and lack of suppression, respectively. In terms of orthographic depth, investigations were administered with Chinese adults (Xu, Kolozsvari, Oostenveld, Leppanen, & Hamalainen, 2019) and typically developing children recently (Xia et al., 2020). In particular, Xia et al. (2020) used Chinese characters and pinyin (a transparent alphabetic coding system that represents the pronunciation of characters, which is taught at the earliest stage of Chinese reading development and used as a scaffold in learning new characters) as experimental materials and observed a significant audiovisual integration effect in the direction of “congruent < incongruent” in the left inferior frontal cortex (IFC) and bilateral STC in processing character-sound associations. Moreover, neural integration in the left IFC in response to character-sound pairs and that in the left STC in response to pinyin-sound pairs were associated with children’s performance in silent reading comprehension that relies on grapho-semantic mapping and oral word reading fluency that relies on grapho-phonological processing, respectively. This pattern is likely to be driven by stimuli’s linguistic properties, including orthographic transparency and involvement of semantics. Using the same experimental design, the current fMRI study aimed to examine whether the neural audiovisual integrations of character-sounds and pinyin-sounds are impaired in Chinese dyslexia and how they associate with different levels of reading abilities.

Notably, while group comparison has been widely used to identify neural deficits in dyslexia, approaches focusing on individual differences also provide invaluable insights. In this case, the brain-behavior correlation is a useful strategy (Jednorog et al., 2015; Pernet, Andersson, Paulesu, & Demonet, 2009), with which two primary patterns can be identified. The first is a universal brain-behavior correlation regardless of reading status (dyslexia vs. control), indicating that the same neural system supports the cognitive processing in both groups. For example, children’s PA is correlated with the microstructural feature of the left arcuate fasciculus, even after controlling group effect (Su et al., 2018; Vandermosten et al., 2012). Alternatively, there could be distinct ways in which reading abilities correlated with brain measures between children with and without dyslexia, indicating dysfunction or compensation (Hoeft et al., 2011; Pernet et al., 2009; Rumsey et al., 1999; Tschentscher, Ruisinger, Blank, Diaz, & von Kriegstein, 2018). For example, while typical readers rely more (higher regional cerebral blood flow) on the left inferior parietal lobule (IPL), higher activation in this area is associated with worse reading performance in dyslexia (Rumsey et al., 1999). However, previous studies commonly conducted correlation analyses while pooling individuals from different groups. Since the participants were selected on purpose, between-group differences could drive the results (Blau et al., 2010). As mentioned above, this issue can be addressed by controlling the effect of group in the statistic model or directly comparing correlations between dyslexia and typical readers.

To summarize, the main aim of this study was to investigate the possible impairments in Chinese children with dyslexia in implicit processing of print-sound associations and related information (e.g., semantics). We asked two specific questions. First, whether the neurofunctional correlates of print-sound integration differ between the dyslexics and controls. Second, whether the relationships between neural integration and reading abilities differ between groups. We adopted a passive fMRI audiovisual paradigm and used characters and pinyin—scripts with contrasting linguistic features—as experimental materials. Both group comparison and individual differences analytic approaches were performed. Based on the prior research in typically developing children (Xia et al., 2020), we predicted a reduced neural integration in Chinese children with dyslexia. In addition, dyslexic children might display atypical brain-behavior correlations or recruit other brain regions to integrate cross-modal information.

## 2 Methods

### 2.1 Participants and behavioral measures

In this study, dyslexia was operationalized by the criteria of having normal intelligence (⩾ 80 on the abbreviated version of the *Chinese Wechsler Intelligence Scale for Children*; Wechsler, 1974) but manifesting reading difficulty (below −1 *SD* of the norm on a standardized reading screening task *Character Recognition*; Xue, Shu, Li, Li, & Tian, 2013). On the other hand, each child in the control group should have normal intelligence (≥ 80) and a score above −0.5 *SD* of the norm on the reading screening task (the aim was to increase the gap in reading skills between groups). In addition, children in both groups should be right-handed (Oldfield, 1971) native speakers of Chinese, with normal hearing and normal or corrected-to-normal vision, and were free from neurological or psychiatric disorders. Finally, only the children that completed all the task fMRI runs, with an overall accuracy equal to or higher than 75% in the in-scanner passive task and less than 25% time-points labeled as outliers (i.e., “bad volume”) in each run (data preprocessing section) were included.

Initially, one hundred children in grades 3-6 (including 45 dyslexia) were recruited from local elementary schools. According to the inclusion criteria described above, 23 dyslexic children (10 girls; age 111-144 months, *M* [*SD*] = 122 [10]) were included in the final analysis. Twenty-one dyslexic children were excluded due to uncompleted MRI data collection (n = 9), severe head motion artifacts (n = 9), or poor in-scanner performance (n = 3). One child with a history of dyslexia diagnosis but performed normally in the character recognition task at the time of data collection was also excluded (this child had received an intensive behavior intervention program). The controls were chosen to match the dyslexic group on grade, age, and sex. The final control group consisted of 22 typically developing children with qualified neuroimaging data (12 girls; age 118-140 months, *M* [*SD*] = 127 [6]).

Each child received a battery of behavioral tests on reading and cognitive-linguistic skills individually in a silent room on the same day of the MRI session. The reading measurements contained: (1) an untimed character naming task (*Character Recognition*) to estimate the number of characters children had conquered (Xue et al., 2013); (2) an oral word reading task (*Word List Reading*) to measure how fast the participant accurately retrieved phonological representations from visually presented high-frequency two-character words (Zhang et al., 2012); and (3) a timed comprehension task (*Silent Reading Comprehension*) to assess the proficiency of meaning access and semantic judgment (Lei et al., 2011). In addition, PA, RAN, and MA—the three most critical cognitive-linguistic skills in Chinese reading acquisition—were measured by *Phoneme Deletion* (Li, Shu, McBride-Chang, Liu, & Peng, 2012), *Digit RAN* (Liu et al., 2017), and *Morphological Production* (Shu, McBride-Chang, Wu, & Liu, 2006).

Written informed consents were obtained from all the children and their guardians after a detailed explanation of the objectives and procedure of the study. After the experiment, each child received a book and a set of stationery as compensation. This study was approved by the Institutional Review Board of the State Key Laboratory of Cognitive Neuroscience and Learning at Beijing Normal University. The data collection was conducted in 2018.

### 2.2 Experimental design

This study adopted a passive audiovisual paradigm that has been widely used in the previous fMRI studies investigating the neural basis of letter-sound integration (**Figure 1**). The details about the stimuli and procedure can be found in Xia et al. (2020). In brief, 56 pictographic characters with high-frequency (*M* ± *SD*= 929 ± 1486 per million; Chinese Single-Character Word Database; https://blclab.org/pyscholinguistic-norms-database/; Liu et al., 2007) and are frequently used as radicals in phonograms were selected. These characters are visually simple (number of strokes: *M* = 4.34, *Range* = 1-9), learned early (age of acquisition: *M*= 3 years, *Range* = 2-5 years), and with high rating scores (7 as the highest value) on concreteness (5.76 ± 1.19) and imageability (5.96 ± 1.00). The pinyin spellings of these characters were used as the visual stimuli in the pinyin experiment. The auditory stimuli (duration: *M* ± *SD* = 476.3 ± 87.5 ms) were the characters’ sounds (i.e., syllables). A native Chinese male recorded the audio files with a sampling rate of 44.1 kHz and 16-bit quantization. The sound files were then normalized to 85 dB and bandpass (100-4000 Hz) filtered with Au dacity (https://www.audacityteam.org/).

**Figure 1.**
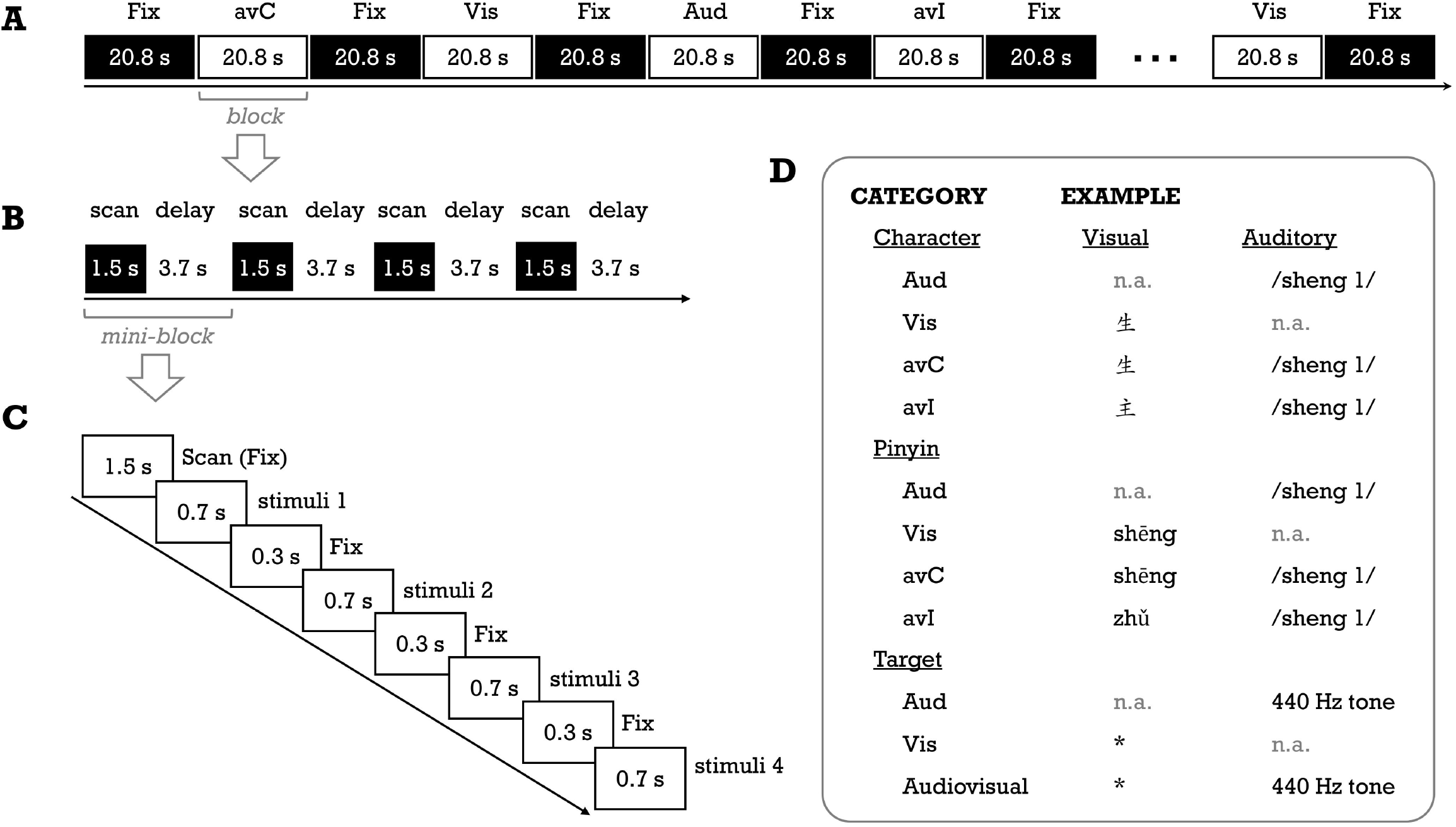
Schematic description of the fMRI experimental procedure (A, B, C for a run, a block, and a mini-block, respectively) and stimuli examples (D). Abbreviations: Aud = auditory, avC = audiovisual congruent, avI = audiovisual incongruent, Fix = fixation, Vis = visual.

The study consisted of 4 task fMRI runs, with the first and second runs for the pinyin experiment and the third and fourth runs for the character experiment. We used the fixed order to prevent priming from characters on visually presented pinyin stimuli. Two unimodal (auditory [Aud]; visual [Vis]) and two cross-modal conditions (congruent [avC]; incongruent [avI]) were created for each experiment (**Figure 1**). In this study, we focused on activation differences between the congruent against incongruent conditions—the neural integration. A block design was used to deliver stimuli. There were 8 task blocks (duration = 20.8 s; 2 blocks for each condition) interleaved with 9 rest blocks (duration = 20.8 s) in a single run. A task block contained 4 mini-blocks. A 1.5 s period was used to collect a whole-brain volume within each mini-block, and a silent period of 3.7 s was used to present stimuli (see “image acquisition” part). The stimuli were presented in white at the center of a black background (*“KaiTi”*; font, 96 pt for characters; *“Century Schoolbook”* font, 90 pt for pinyin). A crosshair was presented at the center of the screen whenever there was no stimulus. To help children keep their attention on the stimuli while avoiding explicit congruency judgment, we used a target detection task (Blau et al., 2010). Specifically, in each task block, two out of 16 experimental stimuli were randomly replaced with the auditory target (440 Hz pure tone), visual (an unpronounceable symbol) target or their combination. The participant was asked to press a button with the right index finger as accurately and quickly as possible whenever the target appeared.

### 2.3 Image acquisition

All brain images were collected at Beijing Normal University Imaging Center for Brain Research using a 3-Tesla Siemens MAGNETOM Trio Tim scanner with a 12-channel head coil. The children first attended a training session to get familiar with the experimental environment and the scanning noise. During the formal scan, foam pads were used to hold their heads secure during scanning to improve image quality. In addition, children could take a break between sequences to reduce the possible fatigue effect. For each participant, two functional runs for the pinyin experiment, one anatomical run for structural images, and two functional runs for the character experiment were administered sequentially. The quality of the brain images was evaluated immediately by a radiologist who was blinded to the details of this study.

The parameters of the functional images (Gradient Echo Planar Imaging [EPI]) were as follows, repetition time, 5.2 s; echo time, 32 ms; acquisition time, 1.5 s; flip angle, 90 degrees; slice thickness, 4.5 mm; interscan gap, 0.675 mm, voxel size, 3.0 × 3.0 × 4.5 mm^3^; 24 slices; 68 volumes). Since this study contained auditory stimuli, to avoid artifacts induced by the noise during scanning, we used a sparse sampling design with 1.5 s for image collection and 3.7 s delay for stimuli presentation (Shah et al., 2000). The parameters of the structural images (Magnetization-Prepared Rapid Acquisition with Gradient Echo [MPRAGE]) were as follows, repetition time, 5.2 s; echo time, 3.39 ms; inversion time, 1.1 s; flip angle, 7 degrees; slice thickness, 1.33 mm; interscan gap, 0 mm; voxel size, 1.3 × 1.0 × 1.3 mm^3^; 144 axial slices).

### 2.4 Data preprocessing

Functional data were analyzed with SPM12 (http://www.fil.ion.ucl). Three dummy scans were added at the beginning of each run to avoid the T1 equilibration effect. No additional volumes were discarded during the preprocessing. The sequence was corrected for head motion. The “bad volumes” were identified with ART-based outlier detection (https://www.nitrc.org/projects/artifact_detect). Considering the age range of participants, a liberal threshold (intensity > 9 *SD*; frame-to-frame head motion > 2 mm) was used. T1 images were segmented and used for transferring the fMRI data from native space to the standard Montreal Neurological Institute (MNI) space. The normalized images were smoothed with an 8-mm Full-Width Half-Maximum Gaussian kernel, and the resulting data were used in the subsequent model estimation. In the 1^st^ level analysis, 4 experimental conditions, 7 head motion parameters (3 for translations, 3 for rotations, and 1 for framewise displacement), and the time point of each “bad volume” were included in the model. The contrast map in which the value of each voxel refers to “avC-avI” (i.e., positive value stands for higher activation in avC than avI; negative value stands for lower activation in avC than avI) was calculated for each child and used in the subsequent analyses.

### 2.5 Statistics

Deficits in reading and reading-related cognitive skills in children with dyslexia were examined at the behavioral level. We first performed descriptive statistics of multiple behavioral tasks in each group and then conducted group comparisons. Next, we calculated correlations between reading and cognitive-linguistic skills within each group and compared them between groups. The Spearman method was used for the variable that was not normally distributed. As for in-scanner performance, a correct response was defined as the button press to the target with a reaction time (RT) ranged from 200 to 2000 ms. Only correct trials were used to calculate the average RT. Finally, the effects of group and run and their interaction were examined on accuracy (ACC) and RT with analysis of variance (ANOVA).

We conducted brain analyses focusing on differences between the two cross-modal conditions for the character and pinyin experiments, respectively, with the same analytic approaches. The nuisance variables of age, sex, and performance IQ were controlled in all the analyses. First, a voxel-wise whole-brain 2 (group: control vs. dyslexic) × 2 (condition: avC vs. avI) ANOVA (i.e., the group comparison approach) was conducted to examine whether the brain regions showing neural integration in children with dyslexia were the same as those in the control group. Significant clusters were identified with the FWE-corrected threshold of *p*-cluster < 0.05 (*p*-voxel < 0.001 for height). These clusters were used as regions-of-interest (ROIs), and post hoc *t*-tests were conducted to interrogate the effects. Complementary ROI analyses were performed to examine correlations between the neural integration with reading and reading-related cognitive-linguistic skills in each group, to identify the reading-related processes involved in audiovisual integration of character/pinyin for typical and dyslexic children, respectively. Once a significant correlation was revealed, we conducted a correlation coefficient comparison between groups.

Next, we used two individual differences approaches to examine the shared and different brain-behavior correlations in children with and without dyslexia in the character and pinyin experiments at the whole-brain level. First, to identify the shared neural basis associated with behavioral performance between groups, we conducted voxel-wise whole-brain regression analyses on the contrast maps of avC against avI across all the participants while controlling for the effects of group, age, sex, and performance IQ. Children’s performance in *Word List Reading* and *Silent Reading Comprehension* tasks were used as regressors in separate models to examine the relationships between neural processing features during print-sound integration with reading abilities that rely more on grapho-phonological mapping and grapho-semantic mapping, respectively. Second, we examined whether the associations between neural integration and reading abilities differ between groups. Same as the previous analysis, we used *Word List Reading* and *Silent Reading Comprehension* as variates of interest in separate models, along with the factor of group and the interaction. In each analysis, an *F*-test was administered with the FWE-corrected threshold of *p*-cluster < 0.05 (*p*-voxel < 0.001 for height), followed by ROI analysis to interrogate the significant effects.

To visualize the results, significant clusters were presented on a FreeSurfer surface template with BrainNet Viewer (Xia, Wang, & He, 2013). Anatomical labeling was performed using the AAL atlas with DPABI (http://rfmri.org/dpabi).

All the behavioral and ROI analyses were administered with SPSS (v24; SPSS Inc., Chicago, IL, USA). Effects were considered significant at*p* < 0.05, and 0.05 < *p* < 0.1 was considered indicative of a trend.

## 3 Results

### 3.1 Behavior

#### 3.1.1 Reading measures and group comparisons

Statistical metrics including *M*, *SD*, *Range,* and result of the Shapiro-Wilk test of each behavioral measurement are presented in Table 1 (see Supplementary Figure 1 for plots). The dyslexic group performed worse in all the reading and reading-related cognitive tasks than the typically developing children (all *p*’s < 0.007). No significant between-group differences were found on age or sex (both *p*’s > 0.05). In addition, the IQs of all the children were within the normal range, while the typical readers had higher scores on both verbal and performance subscales.

**Table 1.** Demographic and behavioral profiles.

| Measures | Typical Readers (n = 22) |  |  | Reading Disorder (n = 23) |  |  | Comparison |  |
| --- | --- | --- | --- | --- | --- | --- | --- | --- |
| | Mean | SD | Range | Mean | SD | Range | *t*/$\chi^2$ | *p*-value |
| Age (month) | 127 | 6 | 118 ~ 140 | 122 † | 10 | 111 ~ 144 | 1.938 | 0.061 |
| Sex (female/male) | 12/10 |  |  | 10/13 |  |  | 0.552 | 0.556 |
| Verbal IQ (standard score) | 109 | 12 | 88 ~ 139 | 97 | 11 | 67 ~ 118 | 3.668 | < 0.001 |
| Performance IQ (standard score) | 115 | 13 | 82 ~ 137 | 105 | 12 | 82 ~ 135 | 2.512 | 0.016 |
| Full-Scale IQ (standard score) | 113 | 11 | 93 ~ 130 | 101 | 9 | 84 ~ 116 | 3.986 | < 0.001 |
| Character Recognition (item) | 122 | 8 | 107 ~ 140 | 86 † | 11 | 53 ~ 99 | 11.830 | < 0.001 |
| Standard score | 1.38 | 0.66 | 0.41 ~ 2.43 | -2.18 | 0.83 | -4.520 ~ -1.020 | 12.701 | < 0.001 |
| Word List Reading (word/minute) | 94 ‡ | 16 | 72 ~ 140 | 63 | 11 | 40 ~ 83 | 7.612 | < 0.001 |
| Standard score | 0.95 † | 1.11 | -0.38 ~ 4.06 | -1.584 | 0.67 | -2.930 ~ -0.130 | 7.278 | < 0.001 |
| Silent Reading Comprehension (character/minute) | 361 ‡ | 104 | 220 ~ 565 | 163 | 56 | 70 ~ 304 | 7.890 | < 0.001 |
| Standard score | 0.98 ‡ | 1.04 | -0.38 ~ 3.12 | -1.083 | 0.57 | -2.250 ~ 0.430 | 7.998 | < 0.001 |
| Phoneme Deletion (item) | 21 † | 4 | 7 ~ 25 | 17 † | 6 | 1 ~ 25 | 2.848 | 0.007 |
| Rapid Naming (second) | 17 | 2 | 11 ~ 20 | 22 | 4 | 15 ~ 30 | -5.481 | < 0.001 |
| Morphological Production (item) | 24 ‡ | 3 | 20 ~ 29 | 22 | 3 | 13 ~ 28 | 2.927 | 0.005 |

Note. † Shapiro-Wilk $p < 0.05$ , ‡ $0.05 < \text{Shapiro-Wilk } p < 0.01$

#### 3.1.2 Correlations between reading and reading-related cognitive skills

Both same and different correlations between reading and cognitive-linguistic skills were observed between children with and without dyslexia. In typical readers, character recognition was significantly correlated with MA (*r*= 0.589, *p*= 0.004) but not PA (*r* = 0.117, *p* = 0.606) or RAN (*r* = −0.301, *p* = 0.174). In dyslexics, however, this ability was associated with PA (*r* = 0.489, *p* = 0.018), but not RAN (*r* = −0.269, *p* = 0.214) or MA (r = 0.008, p = 0.970). The group difference on the correlation coefficients between character recognition and MA was significant (*Z* = 2.09, *p* = 0.037). A similar pattern was found in silent reading comprehension proficiency, where the scores were correlated with MA (*r* = 0.456, *p* = 0.033) in controls but not dyslexia (*r* = −0.012, *p* = 0.957). On the contrast, oral reading fluency was significantly correlated with RAN in both groups (controls: *r* = −0.531, *p* = 0.012; dyslexics: *r* = −0.578, *p* = 0.004).

#### 3.1.3 In-scanner performance

The aim of using a passive target detection task was to ensure that the participant focused their attention on the stimuli delivered via auditory and visual modalities without performing explicit congruency judgment. The results revealed that children in both groups performed the task with high ACC (controls: *M* [SD] = 96.7% [2.9]; dyslexics: *M* [*SD*] = 92.4% [5.0]). In the ANOVA, the main effects of group were significant on both ACC (*p* = 0.001) and RT (*p* = 0.012). The post-hoc analyses showed that the children with dyslexia had lower accuracy and used more time to complete the tasks than the normal controls. The main effect of run was significant on RT (*p* = 0.012; faster as the experiment proceeds) but not on ACC (*p* = 0.645). No significant group × run interaction was observed on either ACC or RT (both *p*’s > 0.05).

### 3.2 fMRI

We used group comparison (ANOVA), and individual differences (brain-behavior correlation) approaches to investigate the impaired neurofunctional features accompanying print-sound integration in Chinese children with dyslexia. In this section, we first present the results of the character experiment, followed by the pinyin experiment.

#### 3.2.1 Character: whole-brain ANOVA and ROI analysis

In the voxel-wise whole-brain ANOVA, the left IFC and STC showed a significant main effect of condition that survived the FWE corrected *p*-cluster < 0.05 (*p*-voxel < 0.001 for height; **Table 2**; **Figure 2A**). The follow-up analysis revealed less activation in the congruent than incongruent conditions in both the controls (LIFC: *t* = 4.361, *p*< 0.001; LSTC: *t* = 3.646, *p* = 0.002) and the dyslexics (LIFC: *t* = 2.346, *p* = 0.028; LSTC: *t* = 3.427, *p* = 0.002; **Figure 2B**). No main effect of group or group × condition interaction survived the whole-brain FWE correction.

**Table 2.** Significant clusters in the voxel-wise whole-brain analyses.

| Experiment and contrast | Label | Brain area | $P_{\text{FWE-corrected}}$ | Size | Peak $F$ | X | Y | Z |
| --- | --- | --- | --- | --- | --- | --- | --- | --- |
| *Character* |  |  |  |  |  |  |  |  |
| Main effect of condition | LSTC | Left middle and superior temporal | 0.007 | 391 | 26.86 | -60 | -32 | 6 |
|  | LIFC | Left inferior frontal gyrus, | 0.024 | 281 | 23.90 | -46 | 16 | 14 |
| *Pinyin* |  |  |  |  |  |  |  |  |
| Group difference in correlation | LOTC | Inferior and middle occipital gyri, | 0.004 | 358 | 29.28 | -44 | -72 | -2 |
| Brain: avC-avI | LSTC | Left middle and superior temporal | 0.039 | 197 | 23.02 | -52 | -44 | 10 |
| Behavior: *Oral reading fluency* | RSTC | Right superior temporal gyrus, | 0.047 | 185 | 38.68 | 62 | -8 | 4 |

*Note.* The significant clusters were identified with the FWE-corrected threshold of $p$ -cluster < 0.05 ( $p$ -voxel < 0.001 for height). Brain area labeling is based on the AAL atlas. Cluster size refers to the number of voxels.

**Figure 2.**
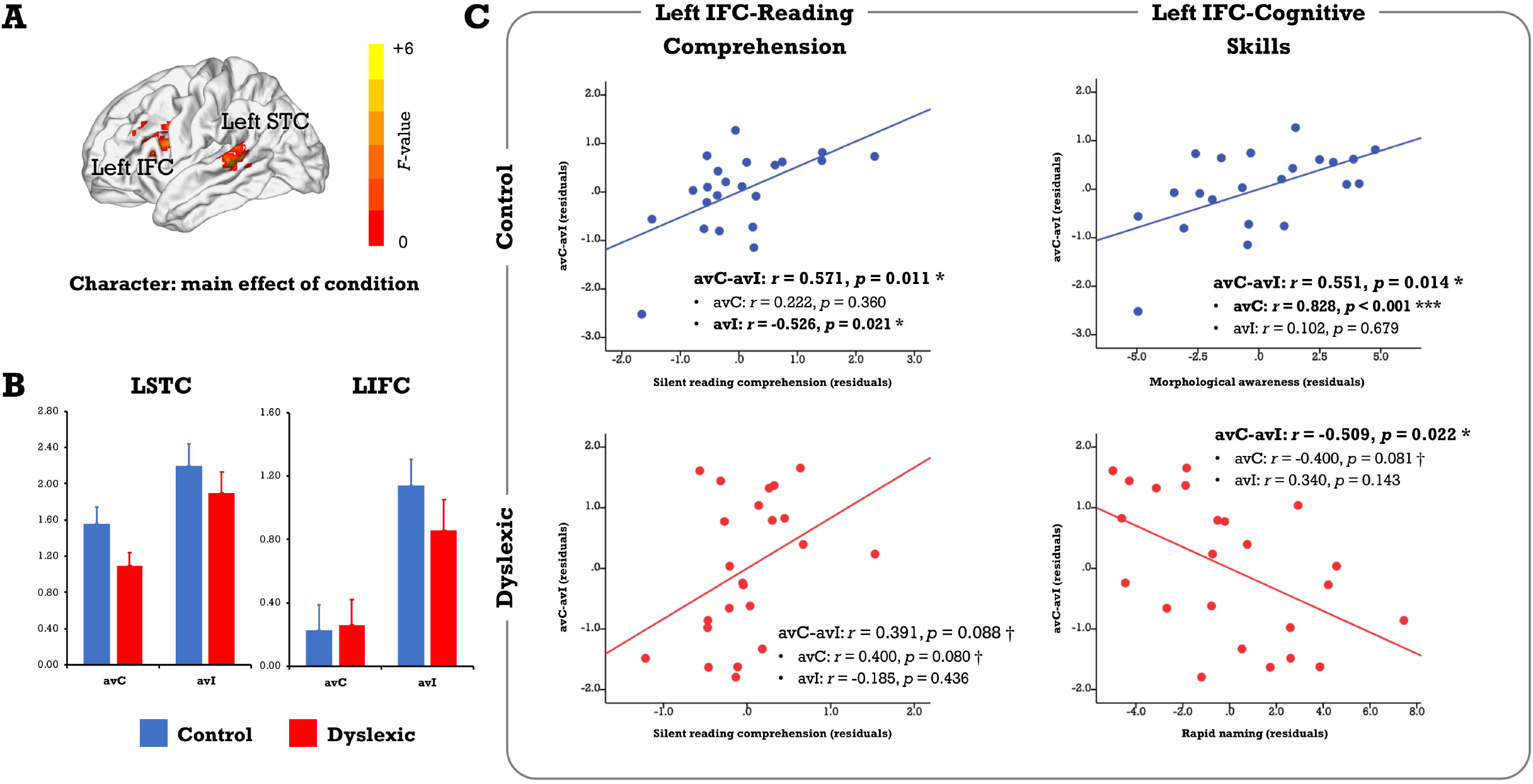
(A) Results of the voxel-wise whole-brain ANOVA on the character conditions. Only clusters showing a significant main effect of condition were identified at the FWE corrected threshold of *p*-cluster < 0.05 (*p*-voxel < 0.001 for height). No region showed significant main effect of group or group × condition interaction. (B) The bar plots present brain activation in the audiovisual congruent and incongruent conditions for the control and dyslexic groups. (C) The scatterplots display correlations between the neural audiovisual integration effect (avC-avI) in the left IFC with silent reading comprehension proficiency and related cognitive-linguistic skills (morphological awareness, rapid naming). Abbreviations: avC = audiovisual congruent, avI = audiovisual incongruent, IFC = inferior frontal cortex, STC = superior temporal cortex.

In the complementary ROI analysis, we observed similar correlations between the neural integration and reading comprehension proficiency in the left IFC in both groups (controls: *r* = 0.571, *p* = 0.011; dyslexics: *r* = 0.391, *p* = 0.088, marginally significant; **Figure 2C**). However, the relative contribution of each cross-modal condition was different. To be specific, the correlation was driven by the incongruent condition in typical readers (avC: *r* = 0.222, *p* = 0.360; avI: *r* = - 0.526, *p* = 0.021) but was more related to the congruent condition in dyslexia (avC: *r* = 0.400, *p* = 0.080, marginally significant; avI: *r* = −0.185, *p* = 0.436). Between-group difference on the correlation coefficients between reading comprehension proficiency and brain activation in the incongruent condition was significant (*Z* = 2.41, *p* = 0.016). The distinct patterns of the correlations between the neural integration and reading-related cognitive skills further support the idea that different mechanisms underlie the integration-comprehension relationships in the two groups (**Figure 2C**): the neural integration in the left IFC was correlated with MA in the controls (*r* = 0.551, *p* = 0.014) but not in the dyslexics (*r* = −0.025, *p*=0.915). Group difference on the correlation coefficients was significant (*Z* = 2.01, *p* = 0.044). On the other hand, the effect was correlated with RAN in the dyslexics (*r* = −0.509, *p* = 0.022) but not in the controls (*r* = - 0.338, *p* = 0.157).

#### 3.2.2 Character: whole-brain group × behavior interaction

Regarding the individual differences approaches, no cluster survived the FWE-corrected threshold of *p*-cluster < 0.05 (*p*-voxel < 0.001 for height) in the analysis investigating the same relationships between the neural integration and reading abilities across groups or that explored the correlation differences between groups.

#### 3.2.3 Pinyin: whole-brain group comparison

The same analytic approaches were used in the pinyin experiment. In the voxel-wise whole-brain ANOVA, no regions showed significant main effect of group or condition or their interaction at the FWE corrected threshold of *p*-cluster < 0.05 (*p*-voxel < 0.001 for height).

#### 3.2.4 Pinyin: whole-brain group × behavior interaction and ROI analysis

We investigated the neural deficits with the individual differences approaches in a whole-brain fashion. While no region displayed the same brain-behavior correlation across groups, clusters located in the left occipitotemporal cortex (OTC) and bilateral STC showed significant between-group differences in the correlation between the neural integration and oral reading fluency (FWE corrected *p*-cluster < 0.05, *p*-voxel < 0.001 for height; **Table 2**; **Figure 3A**). The subsequent analyses revealed positive brain-reading correlations in typical readers and negative correlations in children with dyslexia (**Figure 3B, C and D; Table 3**). Furthermore, the correlations in the left OTC (avC: *r* = 0.685, *p* = 0.001; avI: *r* = 0.118, *p* = 0.631) and STC (avC: *r* = 0.588, *p* = 0.008; avI: *r* = - 0.014, *p* = 0.956) were driven by the congruent condition in the control group, while oral word reading fluency was not correlated with brain activation of the right STC in either the congruent condition (*r* = 0.240, *p* = 0.321) or incongruent condition (*r* = −0.259, *p* = 0.284). In contrast, in dyslexia, the correlations were driven by the incongruent condition (left OTC: *r* = 0.674; *p* = 0.001; left STC: *r* = 0.445, *p* = 0.049; right STC: *r* = 0.543, *p* = 0.013) but not the congruent condition (left OTC:*r* = 0.026, *p* = 0.913; left STC: *r* = −0.050, *p* = 0.833; right STC: *r* = - 0.181, *p* = 0.444) in all three brain regions. Between-group differences on the correlations of the oral reading fluency and activation in the congruent condition in the left OTC (*Z* = 2.54, *p* = 0.011) and left STC (*Z* = 2.26, *p* = 0.024), and on the correlations of the oral reading fluency and activation in the in incongruent condition in the left OTC (*Z* = −2.18, *p* = 0.029) and right STC (Z = −2.73, *p* = 0.006) were significant.

**Figure 3.**
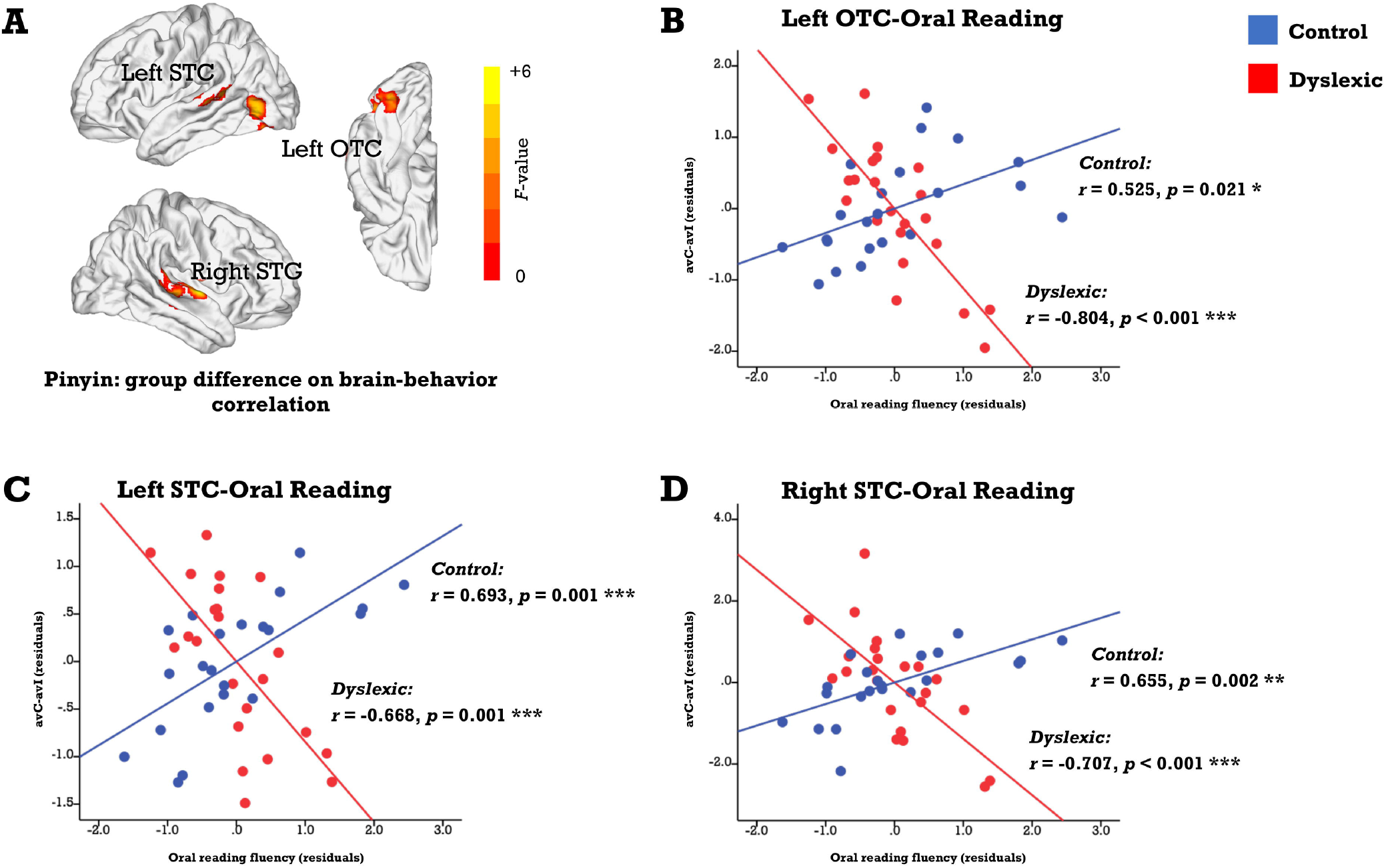
(A) Brain map of regions showing significant between-group differences in the correlation between the audiovisual integration effect and oral reading fluency (FWE corrected *p*-cluster < 0.05, *p*-voxel < 0.001 for height). The scatterplots presents correlations in each group (blue: control; red: dyslexic) in the left OTC (B), STC (C), and right STC (D). Abbreviations: avC = audiovisual congruent, avI = audiovisual incongruent, OTC = occipito-temporal cortex, STC = superior temporal cortex.

**Table 3.** Brain-behavior correlations in the significant clusters in the voxel-wise whole-brain regression analyses.

| Behavior: | Group | Integration Effect (avC-avI) |  | avC |  | avI |  |
| --- | --- | --- | --- | --- | --- | --- | --- |
| *Oral reading fluency* |  | *r*-value | *p*-value | *r*-value | *p*-value | *r*-value | *p*-value |
| L OTC | Control | 0.525 * | 0.021 | 0.685 ** | 0.001 | 0.118 | 0.631 |
|  | Dyslexic | -0.804 *** | < 0.001 | 0.026 | 0.913 | 0.674 ** | 0.001 |
| L STC | Control | 0.693 ** | 0.001 | 0.588 ** | 0.008 | -0.014 | 0.956 |
|  | Dyslexic | -0.668 ** | 0.001 | -0.050 | 0.833 | 0.445 * | 0.049 |
| R STC | Control | 0.655 ** | 0.002 | 0.240 | 0.321 | -0.259 | 0.284 |
|  | Dyslexic | -0.706 *** | < 0.001 | -0.181 | 0.444 | 0.543 * | 0.013 |

*Note.* Age, sex, and performance IQ were controlled statistically. Abbreviations: avC = audiovisual congruent, avI = audiovisual incongruent. \*\*\* $p < 0.001$ , \*\* $p < 0.01$ , \* $p < 0.05$

Regarding correlations between the neural integration and reading-related cognitive-linguistic skills, the left STC was negatively associated with RAN in controls (*r* = −0.479, *p* = 0.038) and showed a trend positively correlated with RAN in dyslexics (*r* = 0.394, *p* = 0.085, marginally significant). The between-group difference was significant (*Z* = −2.93, *p* = 0.003).

## 4 Discussion

This study investigated the neurofunctional features of implicit print-sound integration and their relationships with reading abilities in Chinese children with and without dyslexia. We adopted an fMRI audiovisual paradigm with a passive target detection task, where characters and pinyin—scripts with dramatically different orthographic depths—were used as experimental materials. Of importance, due to the morpho-syllabic nature of characters, semantic information can be automatically activated during character recognition, at least in typical readers. That is, this study enabled us to tap into the three-way relationship between orthography, phonology, and semantics in normal and impaired readers without demanding explicit phonological or semantic processing. Although no between-group differences on the audiovisual integration effect at the brain level were found, the results revealed strikingly atypical correlations between the neural integration of both character-sounds and pinyin-sounds with reading abilities in Chinese children with dyslexia. On the one hand, these anomalies indicate that children with dyslexia rely more on articulatory phonological information during implicit character processing, reflecting a less developed automatic grapho-semantic mapping and integration. On the other hand, it also suggests a malfunctional grapho-phonological mapping in dyslexia and implies that these children may have difficulty in developing the same pinyin processing strategy and transferring it to learning characters as their typically developing peers do.

### 4.1 Left IFC and inefficient grapho-semantic mapping in Chinese dyslexia

First, this study revealed inefficient semantic information access from visual input in Chinese children with dyslexia. We observed slower silent reading comprehension in the dyslexia group. Moreover, while both character recognition accuracy and reading comprehension proficiency were associated with MA in typical readers, children with dyslexia showed significant correlations between reading abilities with other cognitive-linguistic skills instead of MA. Specifically, character recognition was correlated with PA, in line with the previous study with a large independent sample (Song, Zhang, Shu, Su, & McBride, 2020). Regarding silent reading comprehension, although no correlation was significant, there was a trend with RAN, a phonological processing skill that consists of rapid and accurate phonological representation access, retrieval, and articulatory operations. This result is in line with recent studies in which phonological skill contributes to reading more at the early stages of reading acquisition, and the contribution of morphological processing skill increases as children grow (Liu et al., 2017). In addition to the direct effect, MA also mediates the effect of PA on reading (Pan et al., 2016).

Furthermore, we used a passive fMRI audiovisual paradigm to probe the neural bases of processing character-sound associations in Chinese children with and without dyslexia. In terms of brain activation in the left IFC and STC, dyslexic children seem to integrate information from auditory and visual modalities the same way as controls, reflected as a strong audiovisual integration effect in the direction of “congruent < incongruent”. Moreover, the neural integration in the left IFC was associated with silent reading comprehension proficiency, regardless of reading status. However, between-group differences were uncovered in the subsequent ROI analysis. First, while the integration-comprehension association was driven more by the incongruent condition in typical readers, it was driven more by the congruent condition in children with dyslexia, suggesting dyslexic and typical children may use different strategies in processing characters and corresponding sounds, and this difference enlarge with reading abilities increasing within each group. Second, while the audiovisual integration effect in the left IFC was associated with MA in the control group, it was correlated with RAN in dyslexia. These findings indicate that articulatory phonological processing is more likely involved in implicit processing of character-sound pairs in children with dyslexia. In the previous studies investigating neural impairment in Chinese dyslexia, both hypo-and hyper-activation of the left frontal areas were reported (Cao et al., 2016; Liu et al., 2012; Siok et al., 2004). In a meta-analysis, different parts of the left IFC were distinguished based on functionality, where individuals with dyslexia displayed reduced activation in the ventral part associated with semantic processing but increased activation in the dorsal part that was associated with articulatory processing, presumably compensating for their less efficient grapho-semantic route (Hancock, Richlan, & Hoeft, 2017; Richlan, Kronbichler, & Wimmer, 2011). Given that the frontal region is multifunctional (Fedorenko & Blank, 2020; Hagoort, 2014), dyslexia may recruit it in reading-related processing in a different way compared with typical readers.

Of note, the in-scanner task used in this study did not require any sound-semantic or print-semantic processing. Nevertheless, since Chinese has a morpheme-based logographic writing system that involves semantic information even at the character processing level (Guan et al., 2020; Liu et al., 2017; Yang et al., 2013; Zhao et al., 2014), both phonological and semantic information could be accessed effortlessly, at least in typical readers who have received 4-5 years of formal instruction. Thus, it is reasonable to predict that semantic processing skill plays an equal or even more critical role in reading development than phonological processing skill, and its impairments will result in reading difficulties. In line with this hypothesis, previous studies of Chinese demonstrated morphological awareness uniquely predicted reading outcomes and dyslexia status (Pan et al., 2016; Ruan et al., 2018; Song et al., 2020). The current results also showed that while the left IFC was strongly involved in both groups, it was more associated with articulatory phonological processing in dyslexia and semantic-related morphological processing in typical children. The region is close to the one found to underlie morphological processing and show hypo-activation in children with dyslexia during tasks requiring explicit semantic processing (Liu et al., 2013; Zou, Packard, Xia, Liu, & Shu, 2015).

In short, the findings of the character experiment suggest that Chinese children with dyslexia have yet to develop the same brain system for automated semantic access and integration during implicit character/word recognition as that in typically developing children. In contrast, these children are more likely to rely on an articulatory strategy by recruiting the multifunctional frontal area, which may underpin their slow reading comprehension.

### 4.2 Bilateral TPC, left OTC and malfunctioning grapho-phonological mapping in Chinese dyslexia

In addition to the inefficient grapho-semantic mapping, this study also indicates that children with dyslexia may not develop a typical grapho-phonological mapping. At the behavior level, the dyslexic group performed worse in the tasks measuring oral reading fluency, PA, and RAN, in line with previous studies (e.g., Lei et al., 2011). In the pinyin experiment, we observed differential brain-behavior correlations in the classic reading-related areas, including the left OTC and bilateral STC. The morphometric measurements in these regions have also been associated with oral reading fluency in Chinese school-age children (Xia et al., 2018). In particular, the neural integration in the direction of “congruent > incongruent” in response to pinyin-sound pairs was positively associated with oral reading fluency in typical controls: the better the children performed in the oral word reading task, the higher the activation was in the congruent condition than incongruent condition. In contrast, the correlation was negative in dyslexia: children with higher reading fluency showed higher activation in incongruent than congruent conditions. Additionally, the correlations in the control group were driven more by individual differences in brain responses to the congruent stimuli. In contrast, the correlations in the dyslexic group were driven more by the incongruent condition. These findings suggest that while the same brain regions were recruited for both groups’ implicit audiovisual integration of pinyin, children with dyslexia may use them differently.

The OTC and STC in the left hemisphere have been regarded as critical nodes in the classic reading network. Deficits in these areas have also been repeatedly reported in dyslexia (Richlan, Kronbichler, & Wimmer, 2009; Richlan et al., 2011). On the one hand, the left OTC has been considered the interface for initially integrating orthographic, phonological, and semantic information (Price & Devlin, 2011). In addition, the left OTC contains a specific portion in the fusiform gyrus named Visual Word Form Area that has been found to respond specifically to word and word-like stimuli. The left STC, on the other hand, is a central area that represents phonological information (Boets et al., 2013; Glezer et al., 2016), including lexical tone—the supramarginal phoneme in tonal languages such as Chinese (Si, Zhou, & Hong, 2017; L. Zhang et al., 2011). The left STC is functionally and structurally connected to the left OTC, which can be shaped by learning grapho-phonological mappings (Stevens, Kravitz, Peng, Tessler, & Martin, 2017; Thiebaut de Schotten, Cohen, Amemiya, Braga, & Dehaene, 2014). In the current study, besides the left-hemispheric regions, the right STC also showed significant group differences in brain-behavior relationships. Although this region has been less frequently reported in previous studies in alphabetic languages, it subserves lexical tone processing in Chinese (Liang & Du, 2018; Si et al., 2017; Zhang et al., 2011). In addition, the cortical thickness of this area is also associated with oral reading fluency in typically developing Chinese children (Xia et al., 2018).

The differential relationship between the audiovisual integration effect during pinyin-sound processing and oral word reading fluency in dyslexia can be interpreted in at least two ways. First, suppose pinyin processing skill is a continuum, and dyslexia represents the lower end. In that case, the current finding then hints at the expansion and renormalization hypothesis of brain plasticity associated with skill learning (Wenger, Brozzoli, Lindenberger, & Lövdén, 2017). That is, the growth curve of print-sound integration is an inverse U-shape. When the child starts learning pinyin, the brain response to incongruent audiovisual pairs is lower than congruent pairs. With learning, mismatched information extracted from visual and auditory modalities induces higher activation during integration. Then, children turn to focus on overlearned visual scripts by efficiently suppressing attractive auditory information at the highly familiar stage. In this case, activation in the incongruent condition will be suppressed and weaker than the congruent condition. This interpretation is in line with our findings that brain activation in the congruent condition was associated with oral reading in typical readers, whereas incongruent condition was related to oral reading in dyslexia. The alternative explanation is also associated with development but assumes that individuals with dyslexia process pinyin differently from typical controls. In general, typical readers shift from assembled to addressed phonology with reading experience increases (Mei et al., 2014). Pinyin is assembled in nature. But since there are only approximately 400 syllables, it can be expected that typical readers in upper elementary grades who are highly familiar with it could achieve the addressed phonology. Given that children learn Chinese characters as holistic syllable-level units, children who read pinyin with the same addressed phonology may benefit more. In this case, the differential brain-behavior correlations probably reflect the assembled phonology adopted by dyslexic children in processing pinyin-sound pairs. Although this explanation is appealing, conclusions cannot be made without further examination. To date, research on the developmental trajectory of pinyin reading is still lacking. More studies on preliterate and emerging readers with a longitudinal design are needed.

Nonetheless, these findings indicate impaired automatic grapho-phonological mapping in dyslexia from the perspective of individual differences. This anomaly could be underpinned by the altered recruitment of cortical areas such as the left OTC and bilateral STC. As alphabetic languages, learning to read in Chinese requires establishing links between visual forms and linguistic representations (Perfetti & Harris, 2013). Chinese children rely on phonological mediation in reading comprehension at the earliest stages and later gradually shift to rapid grapho-semantic processing with a large amount of practice (Zhou et al., 2017). In this case, deficits in grapho-phonological mapping and corresponding neural basis can impede the development of the ventral pathway for rapid character/word recognition and result in reading problems. Recruiting preliterate children and conducting longitudinal neuroimaging research is necessary to further examine the causal relationship (Nash et al., 2017).

### 4.3 Limitations and future directions

This study has several limitations, and caution should be taken when interpreting the results. First, since we adopted a passive audiovisual integration paradigm here, we could not directly measure the involvement of semantic processing in print-sound processing. Second, we administered the pinyin experiment ahead of the character experiment to reduce the possible prime effect of characters on processing visually presented pinyin stimuli. This may influence brain activation to speech sounds in the character experiment because the same auditory stimuli were used. Third, to have sufficient statistical power, we used the liberal criteria to assess imaging data quality and exclude participants with poor quality data accordingly. Forth, while the overall pattern indicates that children in both groups maintained their attention throughout the experiment, the dyslexia group performed significantly worse. We controlled performance IQ in all the analyses to deal with this issue. The results demonstrated that the main findings of brain-behavior relationships are robust. At last, we did not have enough cases for looking into different subtypes of dyslexia. In the future, studies using multiple experimental designs related to print-sound integration should be conducted with a larger sample size, where a counterbalance design for estimating the order effect, much stricter criteria for controlling MRI data quality, strategies for well-matching on in-scanner performance between groups, and dividing dyslexia into subtypes can be applied.

### 4.4 Conclusion

The present study explored the impaired audiovisual integration of character-sound associations and pinyin-sound associations in Chinese children with dyslexia at the neurofunctional level. The results revealed that dyslexia manifested an atypical relationship between silent reading comprehension and the neural integration of character-sounds in the left IFC and between oral reading fluency and the neural integration of pinyin-sounds in the left OTC and bilateral STC, providing possible neural substrates underpinning inefficient grapho-semantic mapping and grapho-phonological mapping, respectively. Importantly, the current findings also imply that Chinese children with dyslexia may process pinyin—the alphabetic coding system representing the pronunciation of characters—in a lagged or deviated way, which can further impede the development of the direct route for rapid character/word recognition and semantic access.

## 5 Author Contributions

Conceptualization: ZX, HS, XL; Investigation: ZX, TY, XC, HL, XZ; Formal Analysis: ZX; Data Curation: ZX; Writing – Original Draft Preparation: ZX; Writing – Review & Editing: ZX, TY, XC, FH, HL, XZ, HS, XL; Funding Acquisition: ZX, HS, XL; Supervision: ZX, HS, XL

## 6 Acknowledgment

The authors thank all the participating children, their families, and examiners.

## 9 Data availability statement

The data that support the findings of this study are available from the corresponding authors upon reasonable request.

---

**Supplementary Figure 1** The summary of demographic and behavioral measures is shown as boxplots, with the box indicating the IQR. The whiskers show the range of values within 1.5 × IQR and a horizontal line indicating the median. Individual data are shown as dots. The color coding is indicated in the legend below to the plot. Data visualization was performed with PlotsOfData (https://huygens.science.uva.nl/PlotsOfData/). Abbreviations: IQR = interquartile range.

