## Supplementary figures and images for "Atypical relationships between neurofunctional features of print-sound integration and reading abilities in Chinese children with dyslexia"

### Supplemental Figure 1

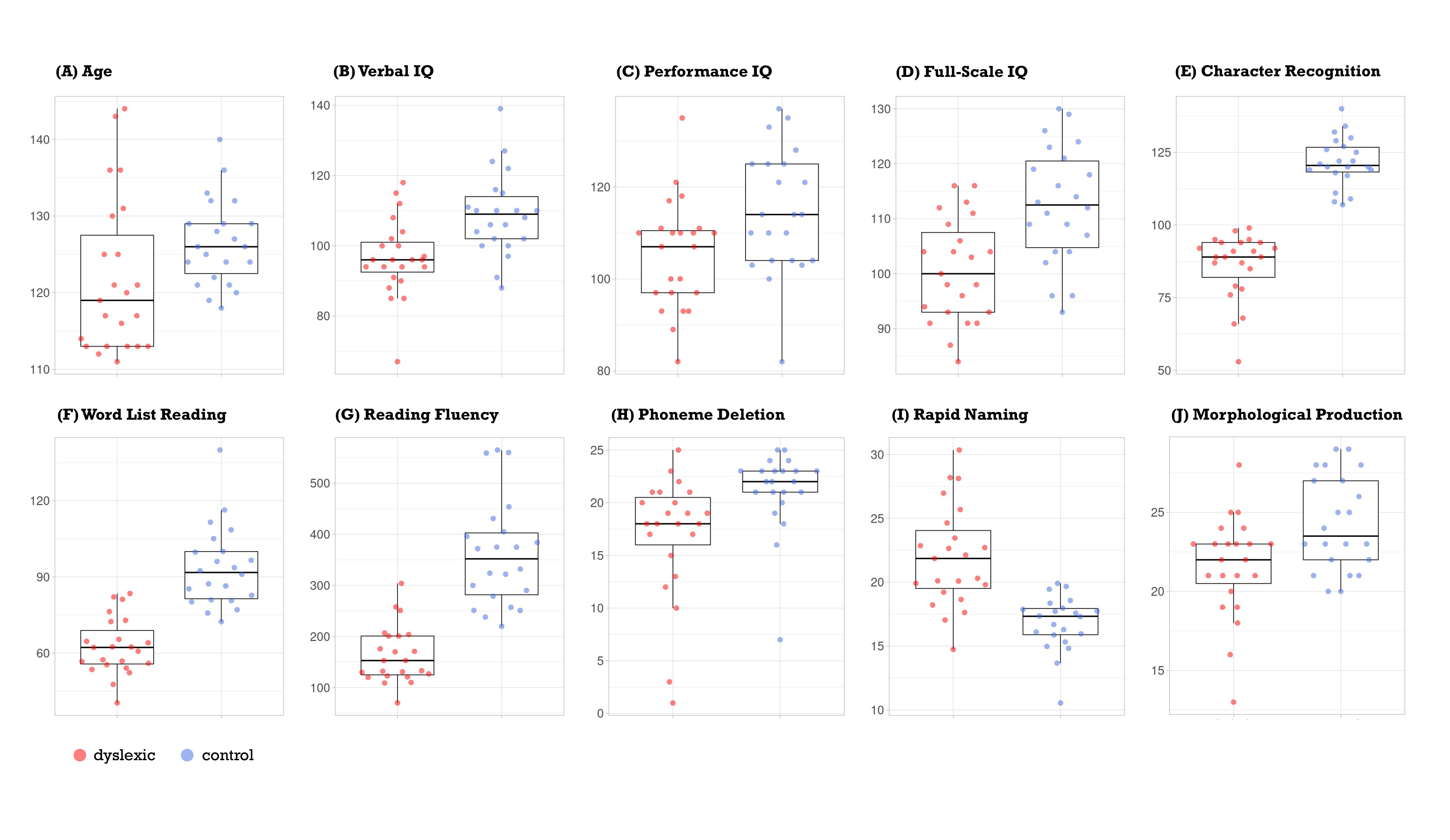
